# Lysine tRNA fragments and miR-194-5p co-regulate hepatic steatosis via β-Klotho and Perilipin 2

**DOI:** 10.1101/2023.09.06.556514

**Authors:** Yonat Tzur, Katarzyna Winek, Nimrod Madrer, Serafima Dubnov, Estelle R Bennett, David S Greenberg, Geula Hanin, Asaad Gammal, Joseph Tam, Isaiah T Arkin, Iddo Paldor, Hermona Soreq

## Abstract

Non-alcoholic fatty liver disease (NAFLD) involves hepatic accumulation of intracellular lipid droplets via incompletely understood processes. Here, we report distinct and cooperative NAFLD roles of LysTTT-5’tRF transfer RNA fragments and microRNA miR-194-5p. Unlike lean animals, dietary-induced NAFLD mice showed hepatic co-declined LysTTT-5’tRF and miR-194-5p levels, restored following hepatic steatosis-suppressing miR-132 antisense oligonucleotide treatment. Moreover, exposing human-derived Hep G2 cells to oleic acid for 7 days co-suppressed miR-194-5p and LysTTT-5’tRF levels while increasing lipid accumulation. Importantly, transfecting fattened cells with a synthetic LysTTT-5’tRF mimic elevated the metabolic regulator β-Klotho mRNA levels while declining triglyceride amounts by 30% within 24 hours. In contradistinction, antisense suppression of miR-194-5p induced accumulation of its novel target, the NAFLD-implicated lipid droplet-coating PLIN2 protein. Further, two out of 15 steatosis-alleviating screened drug repurposing compounds, Danazol and Latanoprost elevated miR-194-5p or LysTTT-5’tRF levels. The different yet complementary roles of miR-194-5p and LysTTT-5’tRF offer new insights into the complex roles of small non-coding RNAs and the multiple pathways involved in NAFLD pathogenesis.

## Introduction

Non-alcoholic fatty liver disease (NAFLD), also referred to as metabolic dysfunction-associated steatotic liver disease (MASLD) is the leading cause of chronic liver disease and one of the most common liver disorders worldwide. NAFLD/MASLD involves hepatic accumulation of protein-coated triglyceride lipid droplets (LDs) [1]. NAFLD begins with simple steatosis, which may progress to non-alcoholic steatohepatitis (NASH) with subsequent higher risk of cirrhosis and hepatocellular carcinoma (HCC) [2]. Often considered as the hepatic manifestation of the metabolic syndrome, NAFLD is associated with obesity, hypertension, dyslipidemia, central adiposity, insulin resistance, and diabetes [3, 4]. Due to its growing prevalence, there is a need to explore the NAFLD underlying mechanisms and seek novel routes for treatment. Since LD accumulation in hepatocytes is the primary characteristic of NAFLD, great efforts have been invested in research aiming to understand LD formation, regulation and contributions to NAFLD progression. LDs are coated by LD-associated proteins such as the perilipins (PLINs), which take part in LD formation, stabilization, and trafficking [5, 6]. Of those, PLIN2 is the major hepatic LD protein. Its levels correlate with steatosis degree [5, 7], and Plin2-deficient mice are NAFLD-resistant and display reduced hepatic triglycerides [8, 9].

NAFLD pathology has been suggested to involve a crosstalk between the liver, gut and adipose tissue [10, 11]. Therefore, understanding the mechanisms leading to NAFLD development and progression is challenging. A significant gut-derived signal that affects adipose tissue and liver response in NAFLD is the fibroblast growth factor (FGF) 15/19 and its receptor system [12]. FGF19 (FGF15 in the mouse), a member of the FGF19 subfamily along with FGF21 and FGF23, is secreted from the intestine and transported to the liver. There, it binds the fibroblast growth factor receptor-4 (FGFR4) assisted by the β-Klotho (KLB) co-receptor, to regulate bile acid (BA) synthesis and homeostasis, which are involved in NAFLD pathogenesis [13, 14]. Thanks to its combining the liver, gut, and adipose tissue contributions [15], and since KLB is a fundamental component that is essential for the complex activation of FGFR and the intracellular downstream responses of FGF signaling [16, 17], KLB and its related proteins as well as the small non-coding RNA regulators (sncRNAs) involved emerged as promising targets for treating NAFLD.

Small non-coding RNAs (sncRNAs) mentioned as causally related to NAFLD initiation and progression include microRNAs (miRs), small (∼22 nucleotides) regulators of gene expression that bind protein-coding mRNA transcripts, silence them post-transcriptionally [18], and regulate numerous biological processes including lipid metabolism, inflammation, and oxidative stress [19–21]. However, while several miRs were implicated in NAFLD pathophysiology [22–24], neither their routes of action nor their complete impact have fully been clarified.

Given the recent re-discovery of transfer RNA fragments (tRFs) as active elements with miR-like capacities [25, 26], we have further pursued potential roles of tRFs in NAFLD. Notably, tRFs are generated by nucleolytic processing of intact tRNAs of nuclear and mitochondrial genome origins, and were recently found to be causally involved in the surveillance of cell proliferation, regulation of gene expression, RNA processing, tumor suppression, and neurodegeneration [27–30]. In ischemic stroke patients, we found tRFs targeted to cholinergic transcripts to replace cholinergic-targeting miRs in nucleated blood cells [31]; and in women living with Alzheimer’s disease, we identified depletion of such tRFs, but not miR stores, to reflect the females-accelerated cognitive decline [32].

To seek miRs and tRFs involvement in NAFLD, we performed small RNA-seq of hepatic tissue from diet-induced obese (DIO) mice before and after antisense oligonucleotide targeting treatment with mmu-miR-132-3p (AM132), which reversed the fatty liver phenotype, compared to matched controls [24]; and used human-derived Hep G2 cells to test the relevance of our findings in the human liver. In both of these systems, we discovered a novel option, whereby complementary yet distinct actions of specific miRs and tRFs emerged as detrimental **joint forces** for NAFLD arrest.

## Results

### LysTTT-originated tRFs and miR-194-5p jointly decline in a murine fatty liver model

Liver tissues of lean regular chow diet (RCD) and DIO mice treated with AM132 or a control antisense oligo targeting hsa-miR-608 (AM608, a primate-specific miR absent in mice, hereafter termed Ctrl) [24] were subjected to small RNA-seq (Figure 1A). Differential expression (DE) analysis revealed altered profiles of both microRNAs (miRs) (Figure 1B) and liver-expressed transfer RNA fragments (tRFs; Figure 1C) in DIO Ctrl mice with the hyperlipidemic phenotype. Principal component analysis (PCA) segregating the levels of distinct treatment-specific patterns revealed specific miRs or tRFs clusters (Figure 1B, 1C) in AM132 compared to Ctrl mice, including 34 upregulated and 47 downregulated miRs, and 13 upregulated and 5 downregulated tRFs (Figure 1D, 1E). Predictably, the AM132 anti-steatotic treatment downregulated miR-132-3p in AM132 mice (log2FoldChange = -12.3 and padj = 4.6e-33), accompanied by miR-122-5p, -148a-5p, -192-5p, let-7g-5p, -21a-5p, -99a-5p, -26a-5p, -194-5p, let-7f-5p and miR-30a-5p, which were the top 10 DE miRs based on their log2Fold change and mean expression levels (count change [33] (Figure 1D, supp. Table 1). Notably, several of those miRs, including miR-122-5p, miR-21, and miR-192 are NAFLD-related [23, 34]. Moreover, KEGG pathway analysis of the predicted targets of all DE miRs [35] showed “*Fatty acid metabolism*” as the top affected pathway (p=1.01E-17), indicating involvement of the identified miRs in the processes leading to liver fat accumulation (Figure 1F, supp. Table 2). In comparison, the 18 DE tRFs originated from nuclear alanine, lysine, and histidine-carrying tRNA genes (supp. Table 3). The top 10 DE molecules by count change measure included 4 Alanine tRNA-derived ones: 3’ tDR-55:76-Ala-CGC-1-M4, 5’ tDR-1:19-Ala-AGC-4-M7, 3’ tDR-55:76-Ala-CGC-3-M2, 5’ tDR-1:25-Ala-AGC-4-M7; 2 Lys-tRNA-derived ones: 5’ tDR-1:18-Lys-TTT-3-M2, 5’ tDR-1:21-Lys-TTT-3-M2, and one each from serine, arginine, histidine and glycine tRNAs; 5’tDR-1:16-Ser-AGA-1-M7, 5’ tDR-1:20-Arg-TCT-1, i-tDR-2:31-His-GTG-1, and 3’ tDR-55:76-Gly-TCC-1-M3 (Figure 1E).

**Figure 1:**
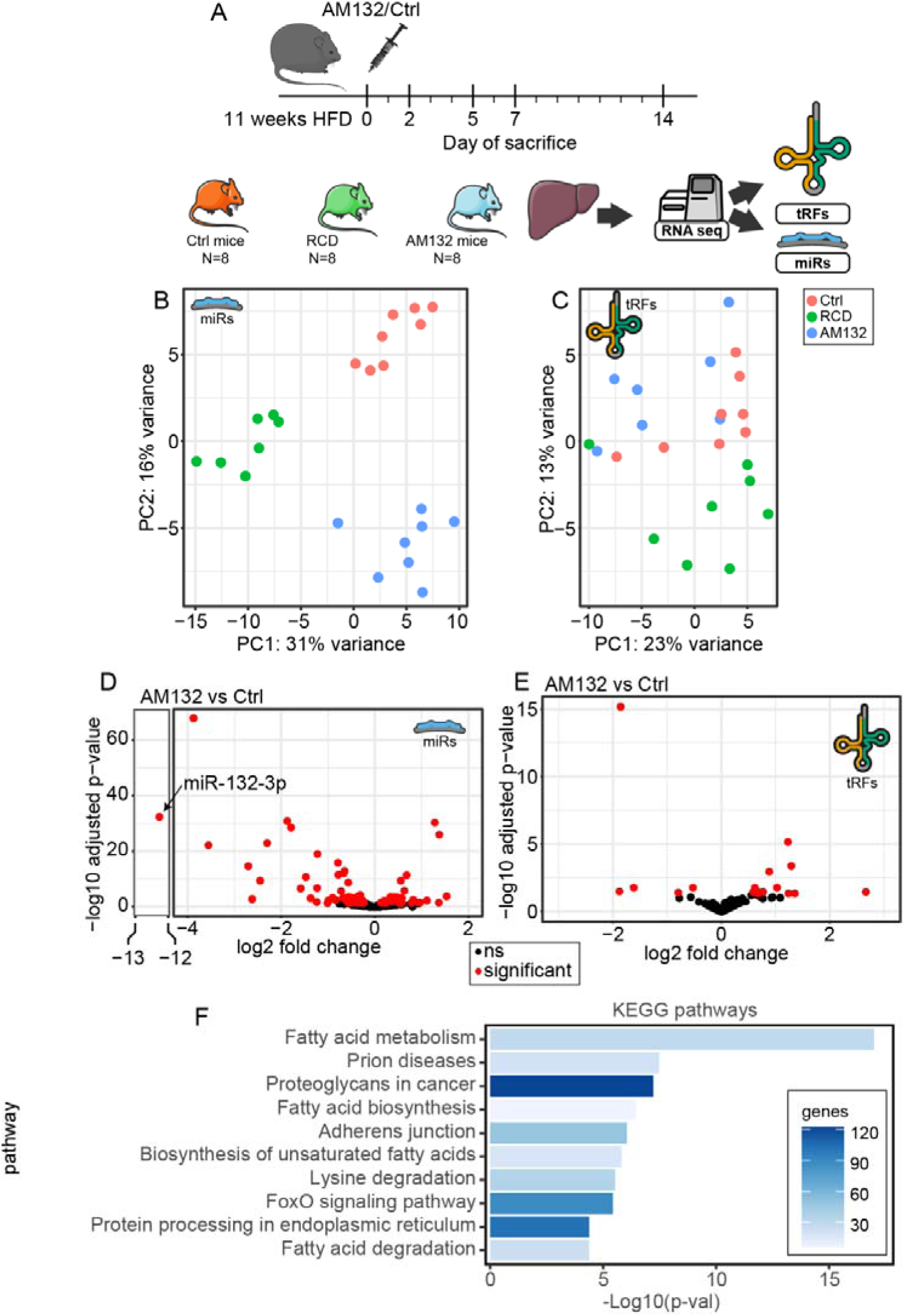
In DIO mice, AM132-reversal of the fatty liver phenotype induced jointly altered miR and tRF profiles. **A.** Experimental design: DIO mice injected with AM132 or AM608 as control (Ctrl) were sacrificed on the stated days, with age-matched RCD mice (n=8 per group). Liver samples were collected and small RNA-seq profiled miRs and tRFs. **B, C.** PCA clustered DE miRs and tRFs according to treatment. **D.** Volcano plot of DE miRs in AM132-treated DIO mice vs control revealed larger fold changes in declined vs elevated miRs. **E.** Volcano plot of DE tRFs in these samples showed mild increases in elevated tRFs. **F.** KEGG pathway analysis of the DE miRs pointed at fatty acid metabolism pathways.

NAFLD’s occurrence and persistence is often linked to altered brain activities, especially in the hypothalamus [36, 37]. Therefore, we tested whether the AM132 treatment also affects hypothalamic small RNAs by subjecting hypothalamic tissues from our NAFLD model mice to small RNA-seq. However, no tRFs, and only two miR changes were observed between the groups (Supp. figure 1), suggesting that the systemic AM132 effects were limited to the liver.

### PLIN2 in Hep G2 and in DIO mice

To investigate the relevance of selected regulatory sncRNAs in the mouse NAFLD model to liver pathophysiology in humans, we exposed human liver-derived hepatocellular carcinoma Hep G2 cells to 0.5 mM oleic acid (OA). Predictably, this triggered dose-dependent lipid accumulation in OA-exposed Hep G2 cells [38], assessed by spectrophotometric quantification of the lipophilic Nile Red-stained intracellular LDs normalized to the number of cells by DAPI-stained nuclei (Figure 2A-B). Importantly, the observed lipid accumulation was accompanied by elevated mRNA and protein levels of the LD-coating perilipin 2 (PLIN2) (Figure 2C, D), which is the most enriched LD-coating protein in the liver, and the levels of which correlate with those of stored triglycerides [5]. Seeking molecular links, we quantified the levels of liver-expressed PLINs in our RCD and AM132 or Ctrl treated DIO mice; notably, Plin2, Plin3, and Plin5 levels were all upregulated in DIO compared to RCD mice, but Plin2 levels alone declined in AM132-treated compared to Ctrl mice (Figure 2E).

**Figure 2:**
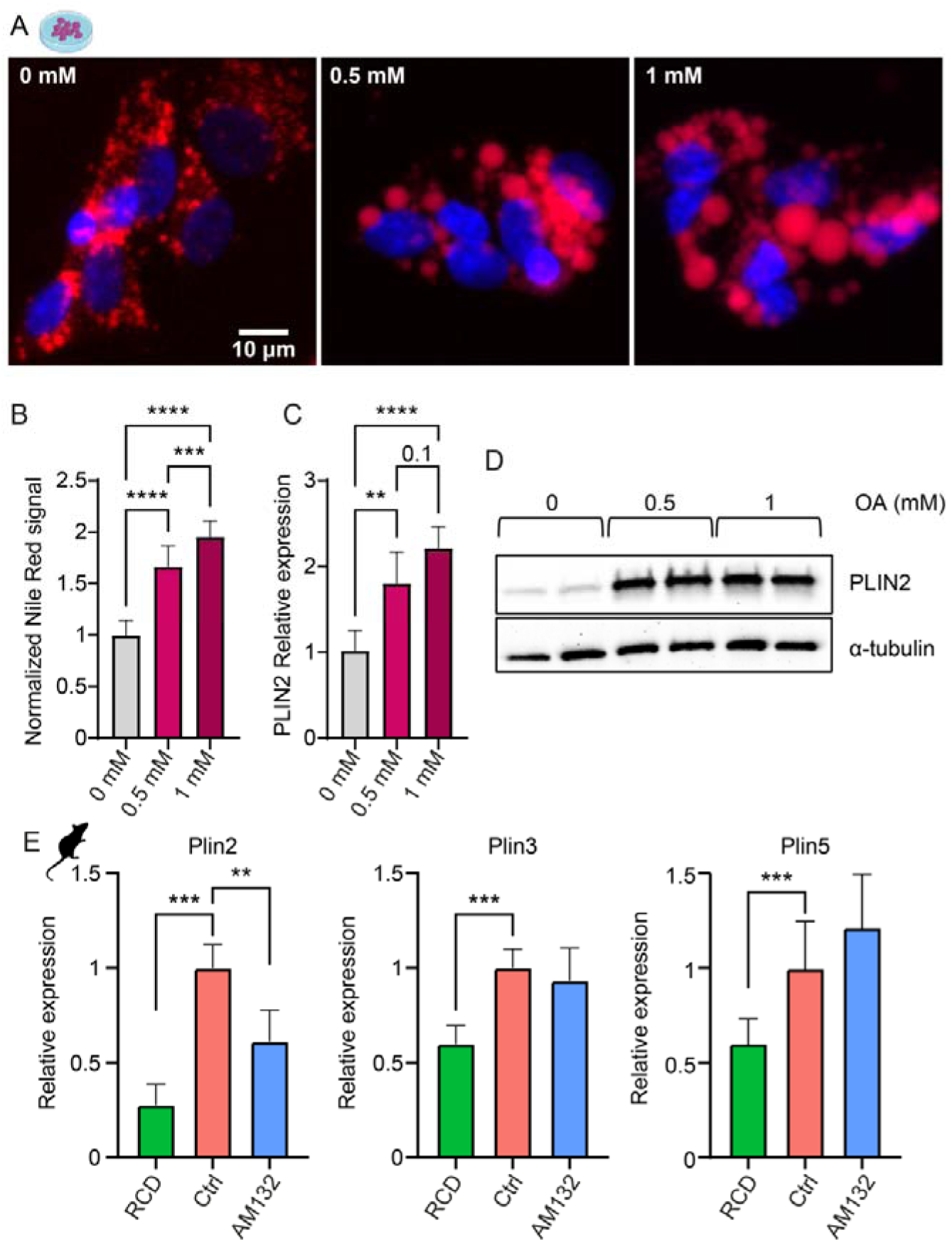
PLIN2 reflects lipid levels dose-dependently in human HepG2 cells and DIO mice. **A.** Fixed Hep G2 cells exposed to 0,0.5, or 1 mM OA for 48 hours showed increasing Nile Red accumulation reflecting TG content (red, Nile Red; blue, DAPI). **B.** Quantification of Nile Red signals in these cells by plate reader measurement, n=3 experiments. **C.** RT-qPCR measurements of PLIN2 mRNA levels normalized to ribosomal protein RPL19 show OA-induced dose-related enhancement of LD formation. **D.** PLIN2 protein levels increase under OA treatment. **E.** AM132 treatment reduced Plin2 (but not Plin3 or Plin5) mRNA levels compared to Ctrl mice; RT-qPCR normalized to b-actin (n=5-8 per group). In panels B, C, E, average ± SD, one-way ANOVA with Tukey’s correction for multiple comparisons, * p <0.05, ** p <0.01, *** p <0.001, ****p<0.0001.

### miR-194-5p directly targets PLIN2

That AM132-treated mice showed lower Plin2 levels than Ctrl mice could either reflect decreased steatosis and reduced presence of lipid droplets-coating proteins at large and/or selective regulation by DE miRs reducing Plin2. To distinguish between these options, we identified the top 10 DE miRs (Figure 1D, supp. Table 1). Among those, the TargetScan target prediction tool [39] identified miR-194-5p as predictably targeting both human and mouse Plin2/PLIN2 (Figure 3A). Correspondingly, RT-qPCR measured miR-194-5p levels were lower in NAFLD-afflicted Ctrl compared to AM132-treated mice (Figure 1D, supp. Table 1, Figure 3B) and in OA-exposed human-derived steatotic Hep G2 cells (∼10% reduction after 48h, and ∼20% after exposure longer than 48h, Figure 3C-E). To further test PLIN2 targeting by miR-194-5p, we knocked-down (KD) miR-194-5p using antisense oligonucleotide (ASO, miR-194-5p, Exiqon) in our Hep G2 cell culture model (Supp. figure 2A). RT-qPCR of miR-194-5p confirmed ∼100% knockdown in both non-steatotic and steatotic Hep G2 cells (Supp. figure 2B-C). Correspondingly, PLIN2 mRNA levels were ∼20% elevated and its protein levels showed 2.5-fold increase following miR-194-5p KD in steatotic cells (Figure 3G-I). To challenge direct PLIN2 regulation by miR-194-5p, we performed dual luciferase reporter assay and co-transfected cells with the 3’UTR of the human PLIN2, and miR-194-5p-/scrambled control sequence-expressing plasmids (Materials and methods). Transfection with miR-194-5p reduced luciferase activity by ∼40%, confirming PLIN2 as a novel target of miR-194-5p (Figure 3J).

**Figure 3:**
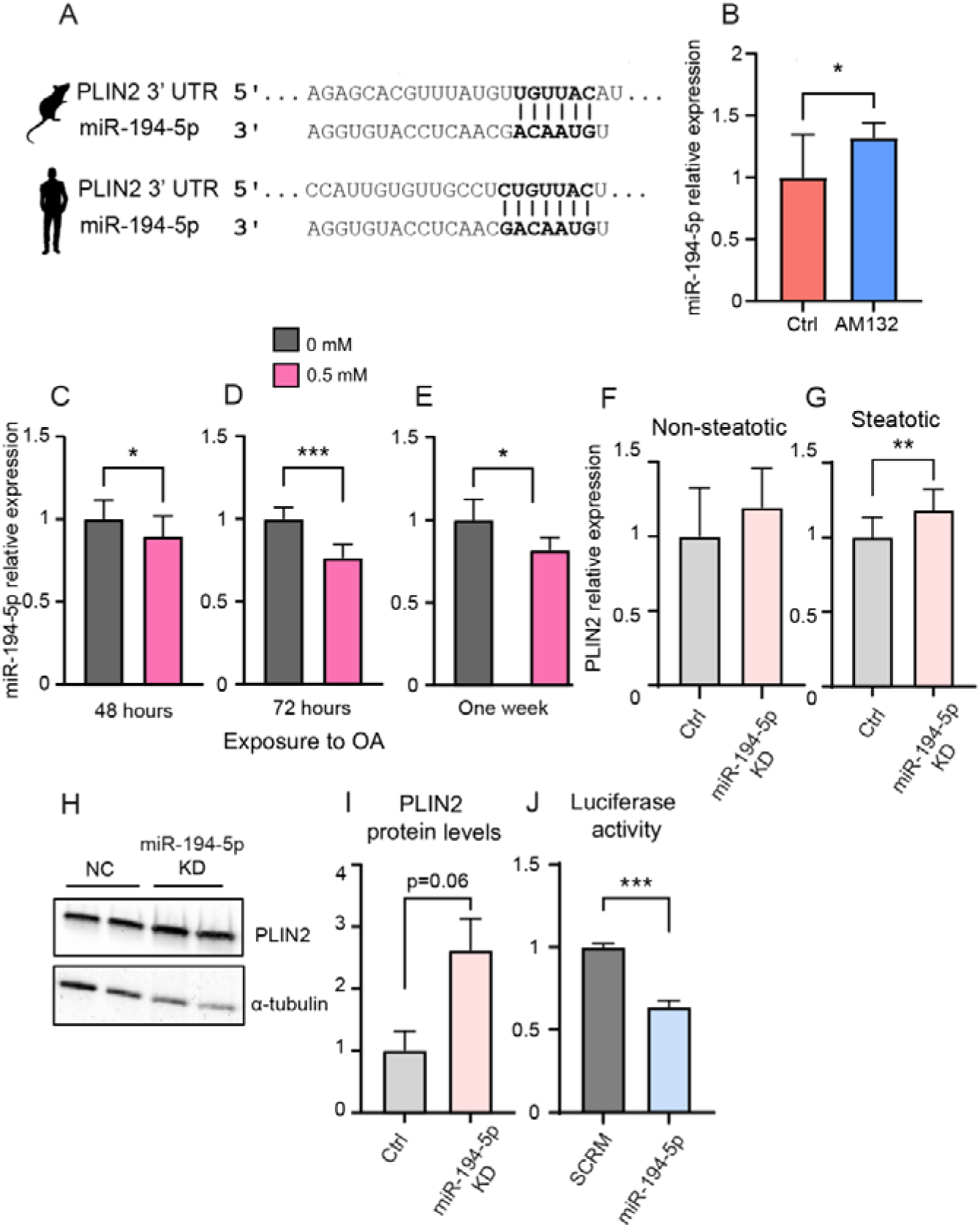
NAFLD-related miR-194-5p decline promotes PLIN2-mediated steatosis. **A.** The predicted binding sites of miR-194-5p in murine and human PLIN2 mRNAs (upper and lower sections). **B.** RT-qPCR validation of RNA-seq presents elevated miR-194-5p levels in AM132 compared to control mice (n=8 per group, RNA-seq in Figure 1D). **C, D, E.** miR-194-5p decline in steatotic Hep G2 cells following 48h, 72h and one week exposure to OA. RT-qPCR measurements normalized to SNORD47, 2-3 experiments each. **F.** Sustained PLIN2 mRNA levels following miR-194-5p KD in non-steatotic cells. **G**. Elevated (∼20%) PLIN2 mRNA levels following KD in steatotic Hep G2 cells. In panels F-G PLIN2 was measured by RT-qPCR and normalized to RPL19, 3 experiments each**. H.** PLIN2 protein increases following miR-194-5p KD in steatotic Hep G2 cells. **I.** Blot quantification of panel H, normalized to a-tubulin. **J.** Decreased luciferase activity in HEK293T cells co-transfected with plasmids expressing human PLIN2 3’UTR and miR-194-5p or a scrambled sequence. In panels B-G, I, J, average ±SD, student’s t-test, * p <0.05, ** p <0.01, *** p <0.001.

### Anti-steatotic AM132 treatment elevates LysTTT-5’tRFs

To explore tRF involvement in NAFLD pathogenesis, we sought liver DE tRFs in the NAFLD-afflicted Ctrl mice which exhibit restored RCD levels following AM132 treatment (Figure 1E, supp. Table 3) [24]. Among the top 10 DE tRFs, two sequence-sharing tRFs originated from LysTTT-tRNA, tDR-1:18-Lys-TTT-3-M2 and tDR-1:21-Lys-TTT-3-M2 (hereafter LysTTT-5’tRFs) fulfilled the criteria of low levels in DIO Ctrl mice, and elevation upon AM132 treatment to the levels observed in RCD mice (Figure 4A). To gain more insight into the kinetics of the LysTTT-5’tRFs expression patterns in NAFLD, we further quantified their levels in liver samples of AM132 and Ctrl mice at 0, 2, 5, 7 and 14 days post-treatment (Figure 4B upper panel) by performing electrophoretic size selection to extract fragments smaller than 50 bases and exclude the amplification of intact tRNAs (Methods; Supplementary Table 5). AM132-treated mice presented progressively reduced liver steatosis, which reached its lowest level at day 7 post-treatment [24]. LysTTT-5’tRFs elevation showed parallel, albeit inverse kinetics initiating on day 5 and 7 post-treatment, reaching peak levels at day 7 and decreasing by day 14 post-treatment (Figure 4B lower panel). The parallel kinetics of elevated LysTTT-5’tRF levels and alleviated steatosis suggested potentially functional involvement of these tRFs in reversing the fatty liver phenotype.

**Figure 4:**
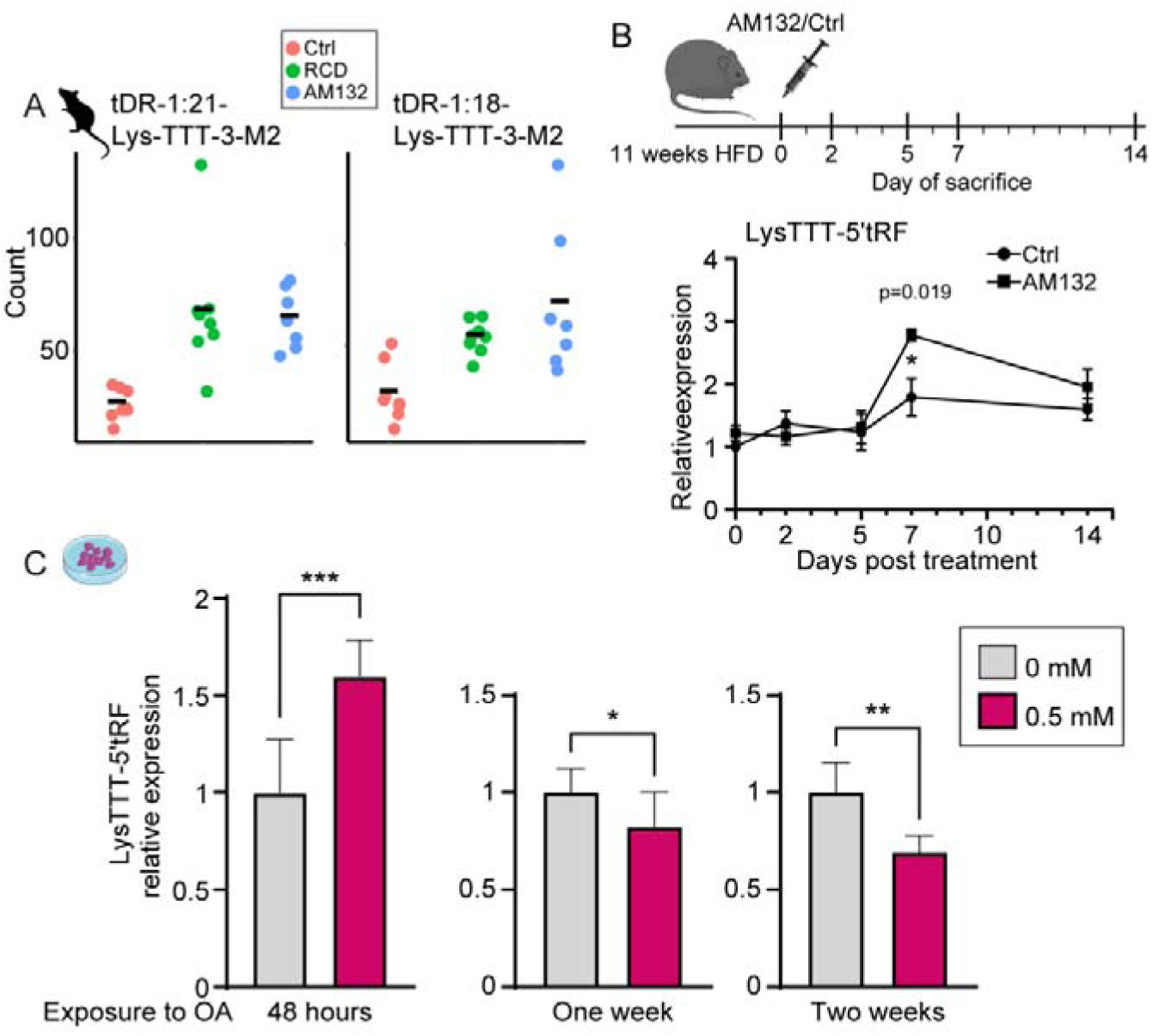
LysTTT-tRFs show NAFLD-related decline in vivo and in-vitro. **A.** The levels of two LysTTT-5’tRFs (tDR-1:21-Lys-TTT-3-M2 and tDR-1:18-Lys-TTT-3-M2) are reduced in NAFLD-afflicted Ctrl mice but restored to lean mice levels under AM132 treatment. **B.** Experimental design (as in Fig.1), with liver samples of AM132-treated and Ctrl mice collected at 0, 2, 5, 7 and 14 post-treatment (n=4 per group). LysTTT-5’tRFs showed highest levels at day 7 after AM132 treatment, when steatosis is lowest. **C.** LysTTT-5’tRF levels increase following acute 48h exposure of Hep G2 cells to OA, and decrease following long-term exposure for one and two weeks. tRF levels were measured by RT-qPCR after size selection and normalized to miR-145-5p and miR-15a-5p, average ±SD, Student’s t-test, * p <0.05, ** p <0.01, *** p <0.001.

To gain more insight into the potential function of LysTTT-5’tRFs, we first tested their levels in our in-vitro Hep G2 steatosis model. Contrasting the decrease observed in the NAFLD-afflicted mice, 48 hours exposure of Hep G2 cells to OA elevated LysTTT-5’tRF levels by 50% (Figure 4C, left). However, since NAFLD is a chronic liver disease where patients are subjected to constant hepatic lipid load, often for years, we suspected that the acute 48 hours OA exposure in our system were insufficient to recapitulate the long-term aspects of NAFLD. To more closely mimic the chronic conditions of exposure to lipid overload, we exposed Hep G2 cells to OA for one and two weeks. Under those conditions, LysTTT-5’tRFs levels were decreased by ∼20% and ∼30% (Figure 3C middle and right graphs), compatible with their reduction in the NAFLD-afflicted mice.

### LysTTT-5’tRF mimic suppresses lipid droplets and elevates **β**-Klotho (KLB) levels in steatotic Hep G2 cells

To pursue causal effects of LysTTT-5’tRFs on triglyceride accumulation, we treated OA-exposed Hep G2 cells to a chemically protected mimic of LysTTT-5’tRF (Figure 5A). Quantifying lipid droplet levels using our computational tool based on microscopic pictures (n=∼150, multiple wells of 3 independent experiments) compared those levels in steatotic cells treated with either a LysTTT-5’tRF mimic or a negative control probe (supp. Materials and Methods), and involved measuring Nile Red staining area (lipid droplets/triglyceride) normalized to the number of nuclei (see supp. Materials and methods and supp. File 1). Within 24 hours, LysTTT-5’tRFs-treated cells showed ∼30% reduced Nile Red staining area compared to steatotic cells treated with an NC probe (NC) (Figure 5B). To re-challenge this observation, we quantified Nile Red staining normalized to nuclear stained DAPI signals by spectrophotometric quantification in independent experiments. Strikingly, Nile Red stained cells treated with the LysTTT-5’tRF mimic showed ∼25% lower signal intensity, reflecting expedite 24 hours-reduced triglyceride levels (Figure 5C). To further seek downstream changes in response to LysTTT-5’tRFs increases, we performed long RNA-seq of steatotic Hep G2 cells treated with LysTTT-5’tRF mimic compared to NC treatment. This analysis revealed 32 DE transcripts, 28 of which upregulated following LysTTT-5’tRF mimic treatment compared to NC (supp. Table 4). Importantly, the transcript with the largest fold change upon LysTTT-5’tRFs mimic treatment was KLB, elevated by ∼25% (log2FoldChange = 0.3, p = 5.69E-06) (Figure 5D). KLB is an essential co-factor of FGFR and is known for its protective role in NAFLD by mediating FGF19/21 signaling to maintain glucose and lipid homeostasis [15]. To verify KLB’s involvement in regulating triglyceride accumulation in our cell culture system, we further measured KLB levels following OA exposure. Notably, KLB levels were reduced in steatotic Hep G2 cells (Figure 5E). Inversely, treatment with a LysTTT-5’tRF mimic elevated KLB levels by ∼35% (Figure 5F), confirming the sequencing result in Figure 5D. Moreover, DIO mice revealed elevated levels of hepatic KLB mRNA levels following AM132 treatment, compatible with our findings in the Hep G2 model (Figure 5G).

**Figure 5:**
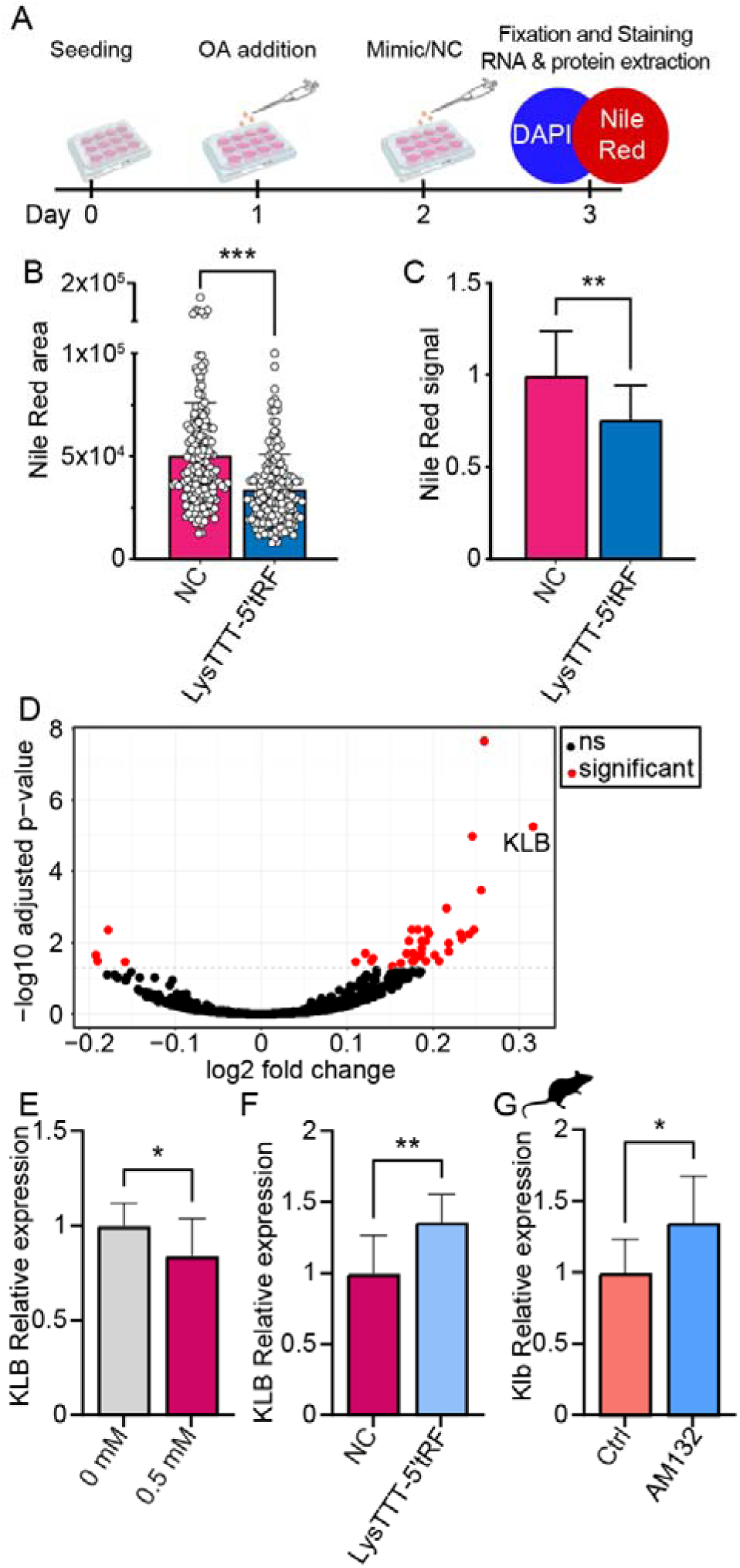
LysTTT-5’tRF suppresses steatosis in Hep G2 cells. **A.** Experimental design: 24 hours after seeding, cells were exposed to 0.5mM OA for 24 hours, then transfected with 100nM of a LysTTT-5’tRF mimic for 24 hours. At 48 hours after OA, 24 hours after mimic treatments cells were fixed, stained and imaged or served for RNA and protein extraction. **B.** TG levels quantification (>150 images), expressed as area of Nile Red signal normalized to the number of nuclei. Note 33% reduction in lipid content following LysTTT-5’tRF mimic treatment. **C.** TG levels by spectrophotometry, expressed as Nile Red normalized to DAPI signal, in 3 independent experiments. Note 25% lower lipid content under LysTTT-5’tRF mimic treatment. **D.** RNA-seq of steatotic Hep G2 cells treated with LysTTT-5’tRF mimic compared to controls, with KLB presenting the most profound fold change in treated cells. **E.** KLB mRNA levels show a decline in steatotic compared to non-steatotic cells. Quantification by RT-qPCR, normalized to RPL19. **F.** Three independent experiments validated the sequencing results shown in panel D and show that KLB mRNA levels were restored in steatotic cells following treatment with a LysTTT-5’tRF mimic. Quantification by RT-qPCR, normalized to RPL19. **G.** Klb mRNA levels were elevated following AM132 treatment compared to Ctrl mice. RT-qPCR, normalized to b-actin (n=7-8 per group). In panels B, C, E-G average ± SD, student’s t-test, * p <0.05, ** p <0.01, *** p <0.001.

### LysTTT-5’tRF and miR-194-5p relate to two different NAFLD-related pathways

To characterize potential pathways/targets which involve LysTTT-5’tRF and miR-194-5p and reduce steatosis, we next performed a drug repurposing screening assay of 1600 drugs that passed phase-I-approved therapeutics (HY-L035, MedChem Express; Monmouth Junction, NJ, USA), identified ones which reduce triglyceride levels, and sought corresponding changes in LysTTT-5’tRF and miR-194-5p levels (Figure 6A). Notably, 15 out of 1600 tested drugs decreased lipid accumulation by over 40% within 24 hours (Figure 6B). Intriguingly, these drugs include anti-infection drugs (Fosamprenavir, Maribavir, Bephenium and Pentoxifylline), anti-inflammation drugs (Diphenylpyraline and Naproxen), and others (Figure 6C). Out of those, **Danazol** (pink) and **Latanoprost** (green) co-elevated the levels of LysTTT-5’’tRF and miR-194-5p. Danazol is a synthetic steroid that suppresses gonadotrophins production and possesses some weak androgenic effects, and Latanoprost (green) is a prostaglandin F2α analog that is used to treat elevated intraocular pressure (IOP). Notably, the remaining drugs showed no effect on the two NAFLED-related sncRNAs (Supp. Figure 3A-B). That only two drugs co-altered the levels of LysTTT-5’tRF and miR-194-5p emphasized the involvement of multiple pathways and various factors in eliciting steatosis, ultimately presenting the complexity of NAFLD pathophysiology and its diverse patterns of cell type specificities.

**Figure 6:**
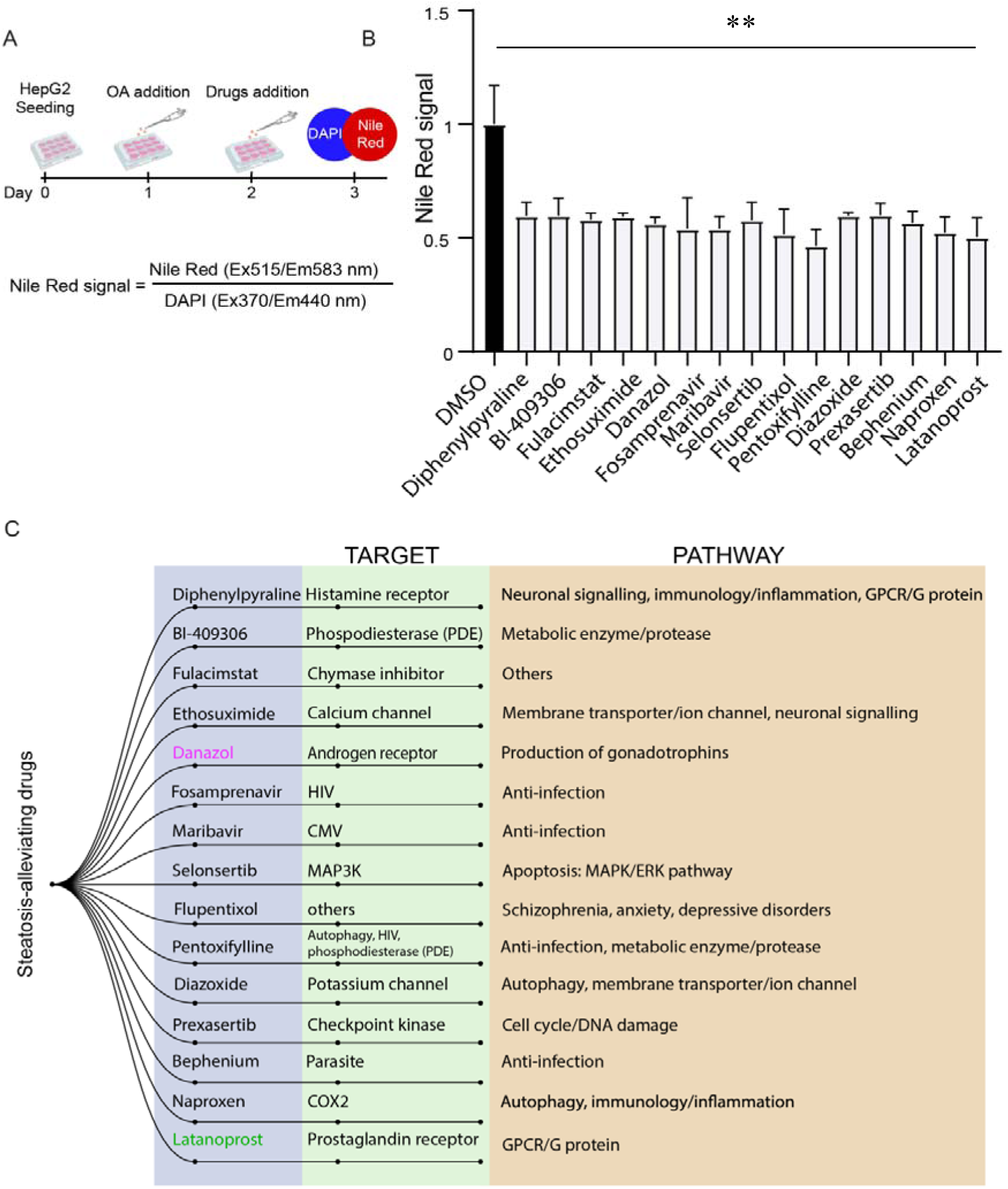
Novel steatosis-alleviating drugs reveal the multilayered complexity of NAFLD. **A.** Experimental design: 24 hours post-seeding, cells were exposed to 0.5mM OA for 24 hours, transfected with 5 µM drugs for 24h, fixed for staining and TG content was quantified by a plate reader, with Nile Red normalized to DAPI signal. **B.** 15 candidate drugs targeting diverse pathways decreased lipid accumulation in steatotic Hep G2 cells by over 40% within 24 hours. **C.** Scheme of the 15 candidate steatosis-alleviating drugs (purple), their known targets and properties (green) and pathways through which they act (brown). Of these, Danazol (pink) and Latanoprost (green) co-elevated the levels of LysTTT-5’’tRF and miR-194-5p. In panel B average ±SD, student’s t-test, * p <0.05, ** p <0.01, *** p <0.001.

## Discussion

Challenging the working hypothesis that NAFLD’s development and progression may involve active role(s) of sncRNAs, we used small RNA-seq to profile the hepatic levels of miRs and tRFs in healthy and fattened mice with or without the steatosis-alleviating AM132 treatment. We identified joint LysTTT-5’tRFs and miR-194-5p declines in NAFLD-afflicted mice and in steatotic HepG2 cells and noted inverse yet transient rescue of their levels following AM132 treatment known to reduce steatosis. Taken together, these findings predicted potential roles of these sncRNAs in human NAFLD.

Unlike the ‘**changing of the guards’** exchange of Cholinergic-targeting miRs to tRFs in blood cells from ischemic stroke patients [31], or the sex- and disease-related loss of tRFs in the brain of women living with Alzheimer’s disease [32], we observed hepatic **‘in tandem’** decline of LysTTT-5’tRFs and miR-194-5p, which may jointly contribute to the steatosis process through complementary impacts on distinct pathways. Supporting human relevance, miR-194-5p decline also occurred in human-originated Hep G2 cells following a short 48h exposure to OA (∼10%), with further reduction after 72h and one week exposure to OA (∼20%). In contrast, LysTTT-5’tRFs levels were first elevated following acute 48h exposure to OA but were reduced following longer 1-2 weeks exposure to OA. Thus, LysTTT-5’tRFs could provide ‘emergency’ control, while miR-194-5p changes presented a longer-term impact. The ‘emergency’ response pattern of LysTTT-5’tRFs is compatible with the fact that no transcription is required to elevate tRF levels, which are processed from already existing tRNAs in the cytoplasm, unlike miRs that first would have to be transcribed in the nucleus and then exported to the cytoplasm [18, 27]. Hence, the observed kinetics is compatible with the fact that tRFs would be more readily available to deal with acute insults. The initial elevation in LysTTT-5’tRF following short exposure to OA is potentially an attempt of the cells to cope with the cellular stress due to the lipid overload exposure, which may lead to enhanced tRNA cleavage and LysTTT-5’tRF production. However, after long exposure to OA, the hepatocytes tRNA reservoir may be depleted, ultimately resulting in decreased levels of LysTTT-5’tRFs and weaker protection from LDs accumulation, and in essence mimicking the female-specific depletion of mitochondrial-originated tRFs from nucleus accumbens [32].

Of particular interest, the identified small RNAs each targets distinct albeit metabolically relevant pathways. Specifically, we uncovered a novel anti-steatotic role of miR-194-5p by identifying the main hepatic LD-coating protein PLIN2 as its direct target. This finding adds to others’ reports of miR-194-5p control over hepatic stellate cell activation which is implicated in the development of liver fibrosis, and of its levels being downregulated in fibrotic liver [40, 41] and in NASH and cirrhosis patients compared to controls, such that it was marked as a potential diagnostic marker for NAFLD [42]. Furthermore, a NAFLD-protective role of miR-194-5p was proposed based on its capacity to decrease the uptake and accumulation of lipids via the Hmgcr and Apoa5 transcripts influencing TG metabolism and fatty acid catabolism; and on its elevated levels in DIO mice treated with green tea over 12 weeks [43]. Adding to its anti-fibrotic effects, our finding of PLIN2 as a miR-194-5p target indicates that the loss of miR-194-5p in NAFLD and fibrosis may further promote steatosis and contribute to NAFLD progression.

Others proposed several tRFs as involved in NAFLD pathophysiology; tRF-3001b may promote NAFLD progression by inhibiting autophagy through targeting the *Prkaa1* gene [44]. Additionally, the levels of three plasma tRFs (tRF-Val-CAC-005, tiRNA-His-GTG-001, and tRF-Ala-CGC-006) elevated in NAFLD patients and mouse models reflect the degree of liver fibrosis [45]. Our finding of LysTTT-5’tRFs protection from NAFLD by reducing TG accumulation in steatotic human-derived HepG2 cells involved restored levels of KLB, which was marked as a promising target for treating NAFLD based on its role as an important mediator of FGF19 and FGF21 signaling [15]. Notably, the KLB rs17618244 variant increases the risk of ballooning and lobular inflammation in children with NAFLD, such that carriers of this single nucleotide polymorphism present lower levels of hepatic and plasma KLB. Furthermore, KLB downregulation in HepG2 and Huh7 cells induced steatosis and upregulation of pro-inflammatory genes [46], and Klb-deficient mice present steatosis and increased levels of pro-inflammatory cytokines [47].

Whether LysTTT-5’tRFs elevation directly regulates KLB mRNA levels requires further identifying of possible mechanism(s) leading to this effect. Importantly, several tRFs are known to affect mRNA stability by interacting with RNA-binding proteins; in breast cancer cells, certain tRFs derived from Glu-, Asp-, Gly-,Tyr-tRNAs suppress the stability of oncogenic transcripts by interacting with the YBX1 protein and displacing it from the 3’UTRs [48]. Also, 5’-tRF derived from Cys-tRNA binds to nucleolin and induces its oligomerization, which in turn enhances nucleolin-bound mRNAs’ stability [49]. However, whether that is the case for LysTTT-5’tRFs and elevated *KLB* levels requires future research.

Intriguingly, both Danazol and Latanoprost, the two approved drugs exerting joint response of LysTTT-5’tRFs and miR-194-5p, target androgen receptors (Danazol) and FP prostanoid receptor (Latanoprost). However, to the best of our knowledge, neither of those was tested for treating NAFLD. In conclusion, we identified two sncRNAs, LysTTT-5’tRF and miR-194-5p which jointly yet distinctly regulate lipid accumulation in hepatocytes via different mechanisms, and indicated multiple pathways for sncRNAs involvement NAFLD development and progression.

## Materials and Methods

### Mice

Animal studies were approved by the ethics committees of The Hebrew University of Jerusalem (approvals 14-14135-3 and NS-16-148663-3). C57bl/6J mice were fed a regular chow diet (RCD) or a high fat diet (Harlan Teklad, Madison, Wisconsin, USA) for 11 weeks to reach diet-induced obesity (DIO). Injected oligonucleotides were modified by locked nucleic acid protection and complementary to mature miR-132 (AM132, 16-mer) or to mature primate-specific miR-608 (AM608, 15-mer, as a control with no predicted complementary sequences in mice) (LNA, Exiqon, Qiagen). DIO mice were injected intravenously with 3.3 mg/kg oligonucleotide for three consecutive days and were sacrificed 0, 2, 5, 7, and 14 days post-treatment. Liver and hypothalamus samples were collected, snap frozen in liquid nitrogen, and stored at - 80°C. Age matched RCD mice were sacrificed and livers collected as above [24].

### Cell culture and tRF mimic experiments

Human hepatocellular carcinoma Hep G2 cells (HEPG2, ATTC HB-8065^™^) were used in all experiments and were grown under standard conditions (Supp. Materials and Methods). Cells were treated with oleic acid (OA)-rich medium (Supp. Materials and Methods) for 24 hours and transfected with 100 nM LysTTT-5’tRF mimic or negative control oligonucleotide (IDT, Syntezza Bioscience, see Supp. Materials and Methods for sequences) using HiPerFect transfection reagent (Qiagen, 301705). After incubation for an additional 24 hours in the same OA-rich medium cells were harvested for RNA and protein or fixed for triglyceride quantification.

### RNA extraction

RNA was extracted using the miRNeasy Mini Kit (Qiagen, 217004) according to the manufacturer’s protocol, followed by RNA concentration determination (NanoDrop 2000, Thermo Scientific), standard gel electrophoresis for quality assesement, and RIN determination (Bioanalyzer 6000, Agilent).

### RT-qPCR

Synthesis of cDNA and RT-qPCR for mRNAs, miRs, and tRFs were done using Quantabio reagents and mouse or human-specific primers (see Supp. Materials and Methods for reagent details and Supp. Table 5 for primer sequences). The CFX384 Touch Real-Time PCR System (Bio-Rad) was used for quantification and the CFX Maestro software (Bio-Rad v4.1.2433.1219) for analysis. Data is presented as relative expression (ΔΔCt) normalized to housekeeping genes and plotted in GraphPad Prism 8.0 (GraphPad Prism Software).

### Immunobloting

Cells were seeded in 12-well plates and incubated in 0, 0.5 or 1mM OA-rich medium for 24 hours prior to transfection with tRF mimic for 24 hours. Cells were then washed and lysed in RIPA buffer, protein concentrations were determined, and 5 µg/sample were loaded and separated by 4-15% gradient polyacrylamide gels and transferred to nitrocellulose membranes. Proteins were detected with antibodies against PLIN2 (Proteintech, 15294-1-AP), KLB (Abcam, ab106794) and Alpha-Tubulin (Merck, T5168), HRP-conjugated secondary antibodies (Jackson, 111-035-

144 and 115-035-062), then ECL (Cell Signalling Technologies, 12757 or Thermo Fisher Scientific, 34580) using the myECL™ Imager and its software (Thermo Fisher Scientific). (see Supp. Materials and Methods for details).

### Triglyceride quantification

Cells were washed, fixed with 4% PFA, permeabilized, and stained with DAPI (1 µg/mL, Santa Cruz Biotechnology, sc-3598) followed by Nile Red (300 nM, Thermo Fisher Scientific, N1142). A Tecan Spark™ 10M plate reader was used for spectrophotometric quantification, with triglyceride content per well expressed as total Nile Red signal normalized to total DAPI signal. Quantification was also carried out by ImageJ analysis [50] of over 150 confocal microscope images (FLUOVIEW FV10i, Olympus) from three independent experiments, with triglyceride content per image expressed as total area of Nile Red signal normalized to number of nuclei (See Supp. Materials and Methods for method details and analysis parameters). The code is publically available, suitable for LD quantification and easy to adjust and repurpose.

### miR-194-5p KD experiments

Cells were seeded in 12-well plates (150,000 cells/well) and transfected 24 hours later with 100 nM antisense oligonucleotide targeting miR-194-5p or miR-608 as control (miR-608 is not expressed in Hep G2 cells) (Exiqon, Qiagen) using Lipofectamine™ 3000 Transfection Reagent (Thermo Fisher Scientific, L3000008). 48 hours later, cells were harvested. For RNA extraction, cells were collected in QIAzol with RNA extraction as above. For protein extraction, cells were washed in PBS and lysed in 140 µl of RIPA buffer (see Supp. Materials and Methods) containing Protease Inhibitor Cocktail and Phosphatase Inhibitor Cocktail (1:100; Cell Signalling Technology, 5871 and 5870). Cells were incubated for 20 minutes on ice, collected, and centrifuged twice (15 minutes, 20,000g, 4°C). Supernatant was transferred to fresh tubes and protein concentration was determined by Lowry assay (Bio-Rad, 5000113).

### Dual luciferase assay

HEK293T cells were seeded in 24-well plates, and 24 hours later co-transfected with 500 ng psiCHECK™-2 Vector (Promega) containing the 3’UTR of human PLIN2 gene downstream to the Renilla luciferase gene, and with 500 ng plasmid expressing either miR-194-5p or a scrambled sequence as control (MR01 backbone, GeneCopoeia). 24 hours later, cells were lysed and assayed using the Dual-Luciferase® Reporter Assay System (Promega, E1910) per the manufacturors instructions, using a Tecan Spark™ 10M plate reader. Results are expressed as ratio of Renilla to firefly luciferase activity for miR-194-5p normalized to scrambled.

### Drug screening assay

1600 drugs of a 2800-compound library (HY-L035, MedChem Express, USA) were screened for identification of novel potential therapeutic targets to alleviate steatosis. Cells were seeded in 96-well plates (20,000 cells/well) and 24 hours later incubated with 0.5 Mm OA. 24 hours later the drugs were introduced at a final concentration of 5 µM with 2.5% DMSO as control. Next, 24 hours later medium was removed and cells were washed and stained for DAPI and Nile Red. TG quantification was performed using a plate reader as describe above and in Supp. Materials and methods.

### Small RNA sequencing and analysis

#### Murine tissues

RNA was extracted as described above from liver (n=8 per group) and hypothalamus (n=12-16 per group) samples of obese mice injected with AM132 or AM608 and lean RCD mice. RIN was determined to be above 8.4 for all samples. Libraries were constructed from 300 ng (liver) or 200 ng (hypothalamus) total RNA (NEBNext Multiplex Small RNA library prep set for Illumina, New England Biolabs, NEB-E7560S) and the small RNA fraction was sequenced on the NextSeq 500 System (Illumina) at the Center for Genomic Technologies Facility, the Hebrew University of Jerusalem.

#### Hep G2 cells

RNA was extracted from steatotic cells treated with LysTTT-5’tRF mimic or NC (two replicates per treatment) as described above and RIN determined to be above 9.5 for all samples. Libraries were constructed from 500 ng total RNA and sequenced as above.

FastQC (http://www.bioinformatics.babraham.ac.uk/projects/fastqc/) was used for quality control and reads were further trimmed and filtered using Flexbar (version 0.11.9 [51]) [2]. Sequences were aligned using miRExpress 2.1.4 [52] (liver) or miRDeep2 [53] (hypothalamus) to miRBase version 21 for micrRNAs and to MINTmap 1.0 for tRFs [54]. Differential expression analysis was performed in R version 4.1.3 using DESeq2 [55]. In liver shrinking log2fold changes was calculated with the “apeglm” algorithm [56] in small RNA expression datasets and “normal” in mimic experiments. Outlier samples in liver were excluded based on PCA plots.

### tRF size selection and qPCR

To validate sequencing results we isolated tRFs from total RNA samples, while excluding whole tRNA molecules, as follows: 1 ug total RNA per sample were separated by PAGE and bands containing RNAs less than 50 nucleotides long were excised from the gel based on two size markers. RNA was eluted from the gel, ethanol-precipitated, and recovered in ddw. cDNA was synthesized from 500 pg/sample and tRF levels were quantified by qPCR as described above. (See Supp. Materials and Methods for detailed protocol).

## Funding and Acknowledgements

The research leading to these results received funding by the Israel Science Foundation, ISF (1016/18; to H. Soreq) and by a joint research support to H. Soreq and I. Paldor from Hebrew University and the Shaarei Zedek Medical center. YT’s and NM’s fellowships were supported by the US friends of the Hebrew University of Jerusalem and The Sephardic Foundation on Aging (New York). KW was a Shimon Peres post-doctoral Fellow at ELSC and SD is an Azrieli PhD fellow. Parts of the figures in this article were created with BioRender.com.

## Data availability statement

The original gene expression data (FASTQ files, metadata and tables of raw counts) from the mice and Hep G2 and all other relevant datasets have been included as supplementary files to the manuscript and the original files and code are available upon request from corresponding author.

## Supplementary material

**Supp. Table 1:**
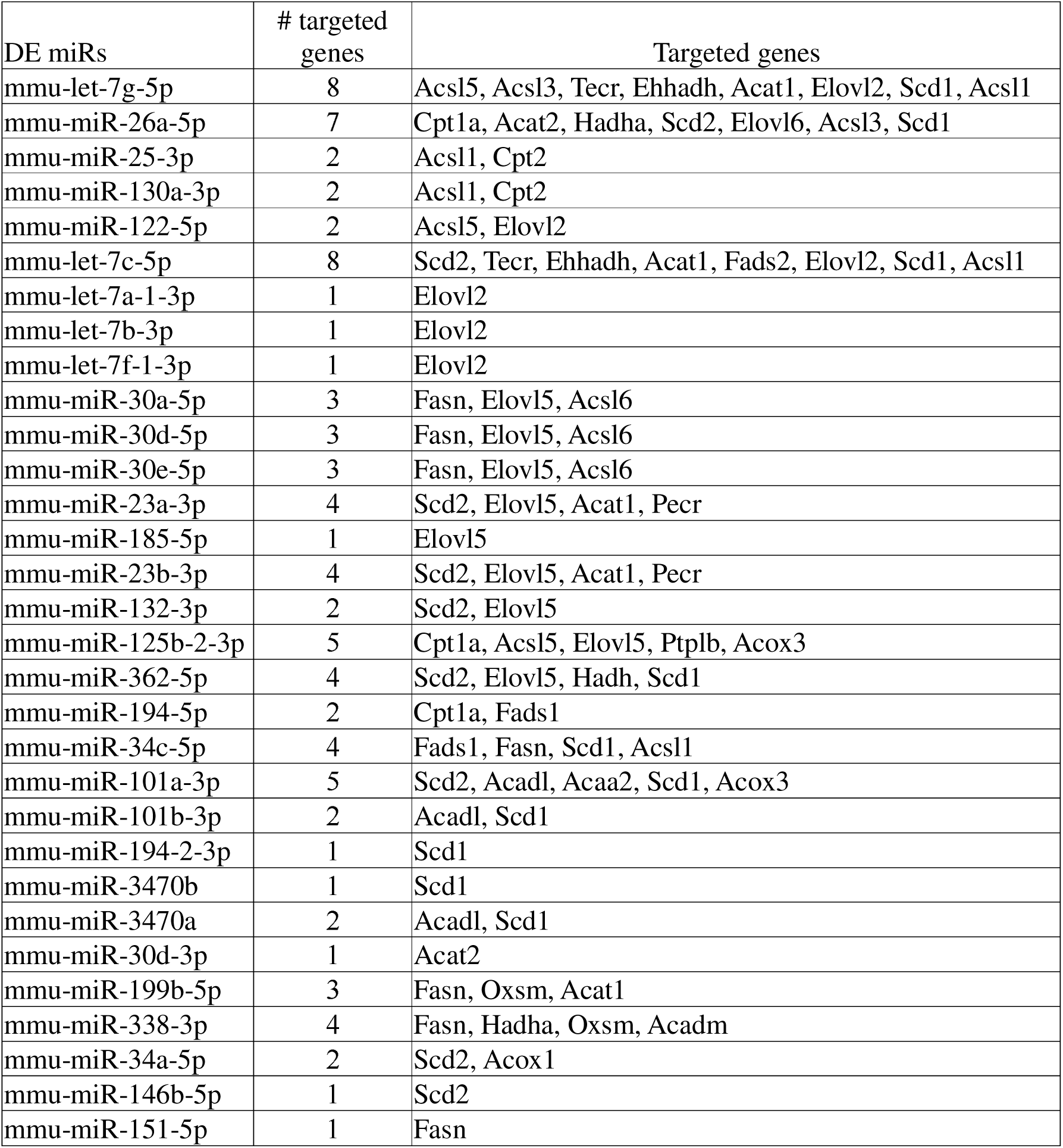
differentially expressed (DE) miRs in AM132 compared to AM608 control mice. **Submitted separately**

**Supp. Table 2:** A list of the DE miRs involved in ’Fatty acid metabolism’ according to KEGG analysis and their targeted metabolic genes.

**Supp. Table 3:** DE tRFs in AM132 compared to AM608 control mice. **Submitted separately**

**Supp. Figure 1:**
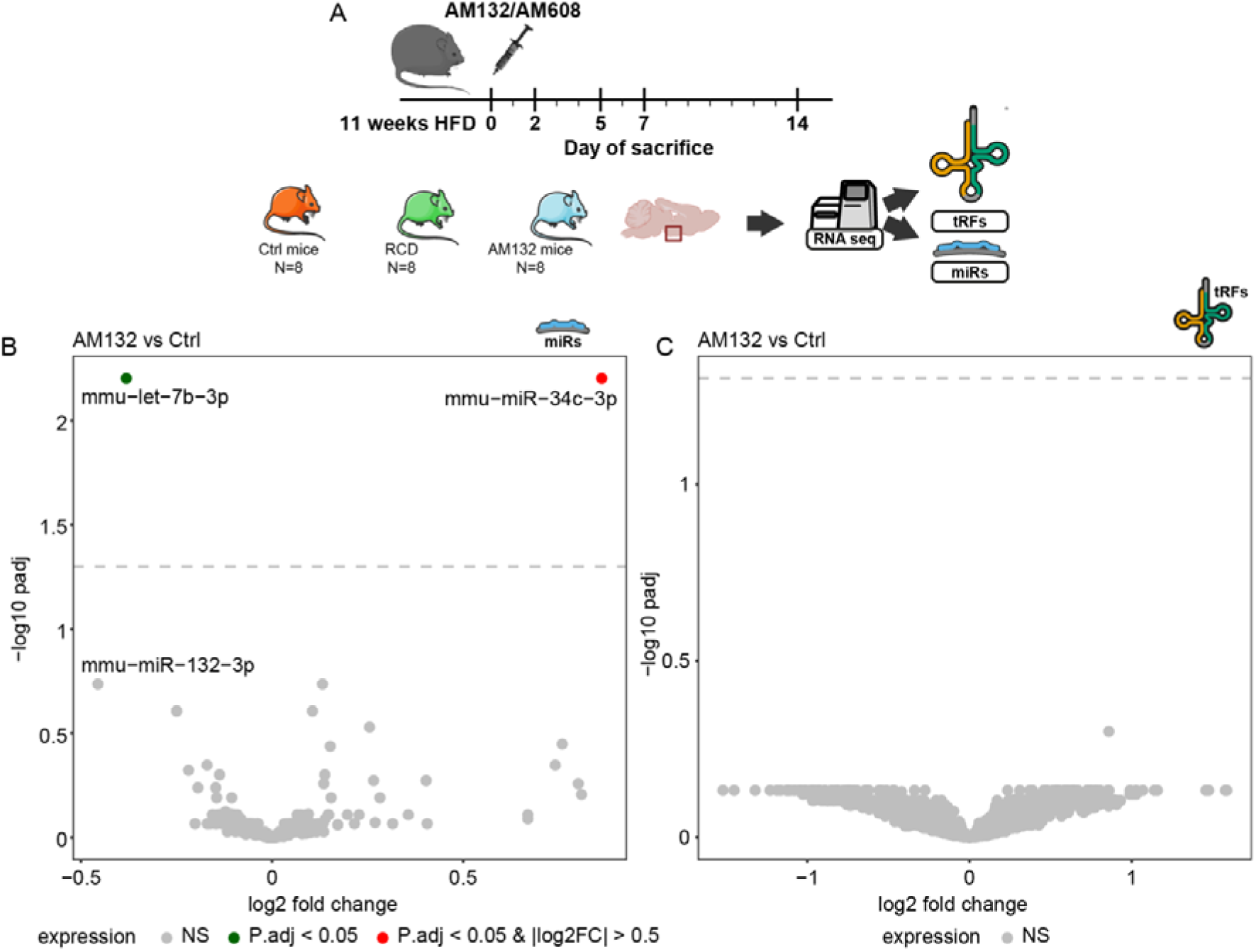
Minor changes in sncRNA levels in hypothalamus suggest local hepatic effect of AM132 treatment. A. Hypothalamus tissues from AM132, Ctrl and lean regular chow diet-fed (RCD) mice served for miR and tRF profiling (N=12-16 per group). B. C. Volcano plots revealing two DE miRs (let-7b-3p; log2FC = -0.38, p = 0.006 and miR-34c-3p; log2FC = 0.86, p = 0.006) and no changes in tRF levels in hypothalamus of AM132-treated mice compared to Ctrl.

**Supp. Figure 2:**
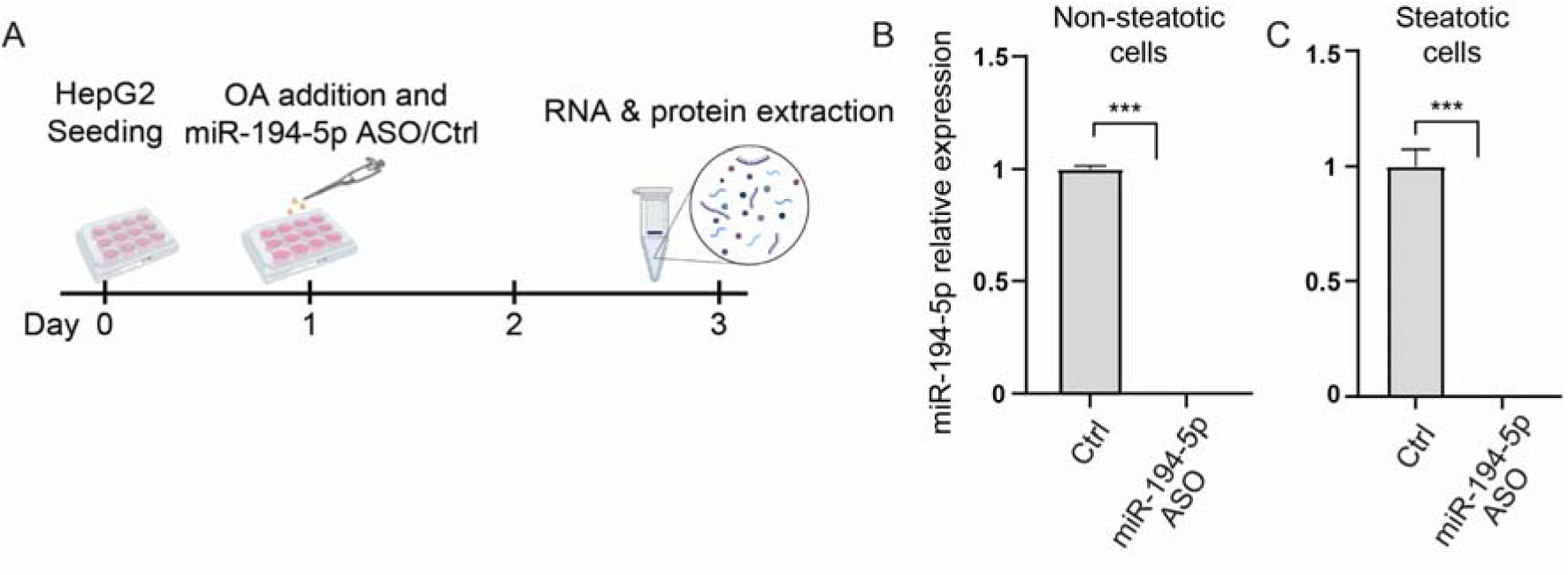
Efficient miR-194-5p depletion following antisense oligo (ASO) treatment in Hep G2 cells. A. Experimental design: 24 hours after seeding, cells were exposed to 0/0.5mM OA and transfected with ASO for miR-194-5p or ASO for miR-608 as control (miR-608 is not expressed in Hep G2 cells, data not shown, Ctrl). 48h after, cells were harvested and RNA and protein were extracted B. miR-194-5p levels in non-steatotic cells, following miR-194-5p ASO treatment compared to control, validating full depletion of the miR. C. The same measurement and depletion are seen in steatotic cells. miR-194-5p was measured by RT-qPCR and normalized to SNORD47. In panels B-C average ±SD, student’s t-test, * p <0.05, ** p <0.01, *** p <0.001.

**Supp. Table 4:** DE transcripts in steatotic Hep G2 cell treated with LysTTT-5’tRF mimic compared to NC probe.

|  | ENSEMBL | gene_name | baseMean | log2FoldChange | padj |
| --- | --- | --- | --- | --- | --- |
| 1 | ENSG00000134962 | KLB | 3055.538 | 0.31611 | 5.69E-06 |
| 2 | ENSG00000152952 | PLOD2 | 15852.9 | 0.259275 | 2.26E-08 |
| 3 | ENSG00000120705 | ETF1 | 4810.298 | 0.255635 | 0.000342 |
| 4 | ENSG00000111328 | CDK2AP1 | 1638.586 | 0.247422 | 0.004342 |
| 5 | ENSG00000104635 | SLC39A14 | 21891.06 | 0.245377 | 1.05E-05 |
| 6 | ENSG00000165959 | CLMN | 1657.653 | 0.241908 | 0.005736 |
| 7 | ENSG00000096088 | PGC | 1567.867 | 0.233415 | 0.007728 |
| 8 | ENSG00000144959 | NCEH1 | 2109.22 | 0.231656 | 0.005496 |
| 9 | ENSG00000113916 | BCL6 | 2373.634 | 0.218186 | 0.017571 |
| 10 | ENSG00000134602 | STK26 | 1787.41 | 0.218054 | 0.010318 |
| 11 | ENSG00000152127 | MGAT5 | 4539.691 | 0.21552 | 0.0011 |
| 12 | ENSG00000224189 | HAGLR | 1509.326 | 0.206936 | 0.032798 |
| 13 | ENSG00000100906 | NFKBIA | 1995.485 | 0.201853 | 0.022175 |
| 14 | ENSG00000188488 | SERPINA5 | 6673.022 | 0.195158 | 0.005496 |
| 15 | ENSG00000071073 | MGAT4A | 5652.846 | 0.193054 | 0.004342 |
| 16 | ENSG00000124839 | RAB17 | 2051.531 | 0.191528 | 0.032798 |
| 17 | ENSG00000152894 | PTPRK | 6599.248 | 0.19133 | 0.008896 |
| 18 | ENSG00000099991 | CABIN1 | 5326.915 | 0.186665 | 0.014542 |
| 19 | ENSG00000115419 | GLS | 4638.781 | 0.186639 | 0.008896 |
| 20 | ENSG00000100644 | HIF1A | 3388.869 | 0.186214 | 0.019808 |
| 21 | ENSG00000147526 | TACC1 | 3420.974 | 0.183759 | 0.024824 |
| 22 | ENSG00000166123 | GPT2 | 8681.11 | 0.182237 | 0.004342 |
| 23 | ENSG00000115902 | SLC1A4 | 3048.872 | 0.177414 | 0.032798 |
| 24 | ENSG00000142192 | APP | 8735.362 | 0.176474 | 0.019808 |
| 25 | ENSG00000129657 | SEC14L1 | 4754.03 | 0.17539 | 0.032798 |
| 26 | ENSG00000089220 | PEBP1 | 10834.18 | 0.174966 | 0.004342 |
| 27 | ENSG00000143153 | ATP1B1 | 6292.132 | 0.171732 | 0.008896 |
| 28 | ENSG00000059573 | ALDH18A1 | 6517.973 | 0.168919 | 0.020062 |
| 29 | ENSG00000118985 | ELL2 | 6146.018 | 0.162247 | 0.03755 |
| 30 | ENSG00000005893 | LAMP2 | 4726.232 | 0.152461 | 0.046733 |
| 31 | ENSG00000135048 | CEMIP2 | 13636.54 | 0.130166 | 0.027163 |
| 32 | ENSG00000118785 | SPP1 | 23467.07 | 0.128176 | 0.032798 |
| 33 | ENSG00000189056 | RELN | 29431.14 | 0.121099 | 0.019808 |
| 34 | ENSG00000083857 | FAT1 | 34683.86 | 0.109899 | 0.034307 |
| 35 | ENSG00000106366 | SERPINE1 | 13501.2 | -0.15823 | 0.034509 |
| 36 | ENSG00000167244 | IGF2 | 12249.18 | -0.17798 | 0.004432 |
| 37 | ENSG00000146038 | DCDC2 | 2677.161 | -0.19027 | 0.032798 |
| 38 | ENSG00000165802 | NSMF | 2576.582 | -0.19222 | 0.022175 |

**Supp. Figure 3:**
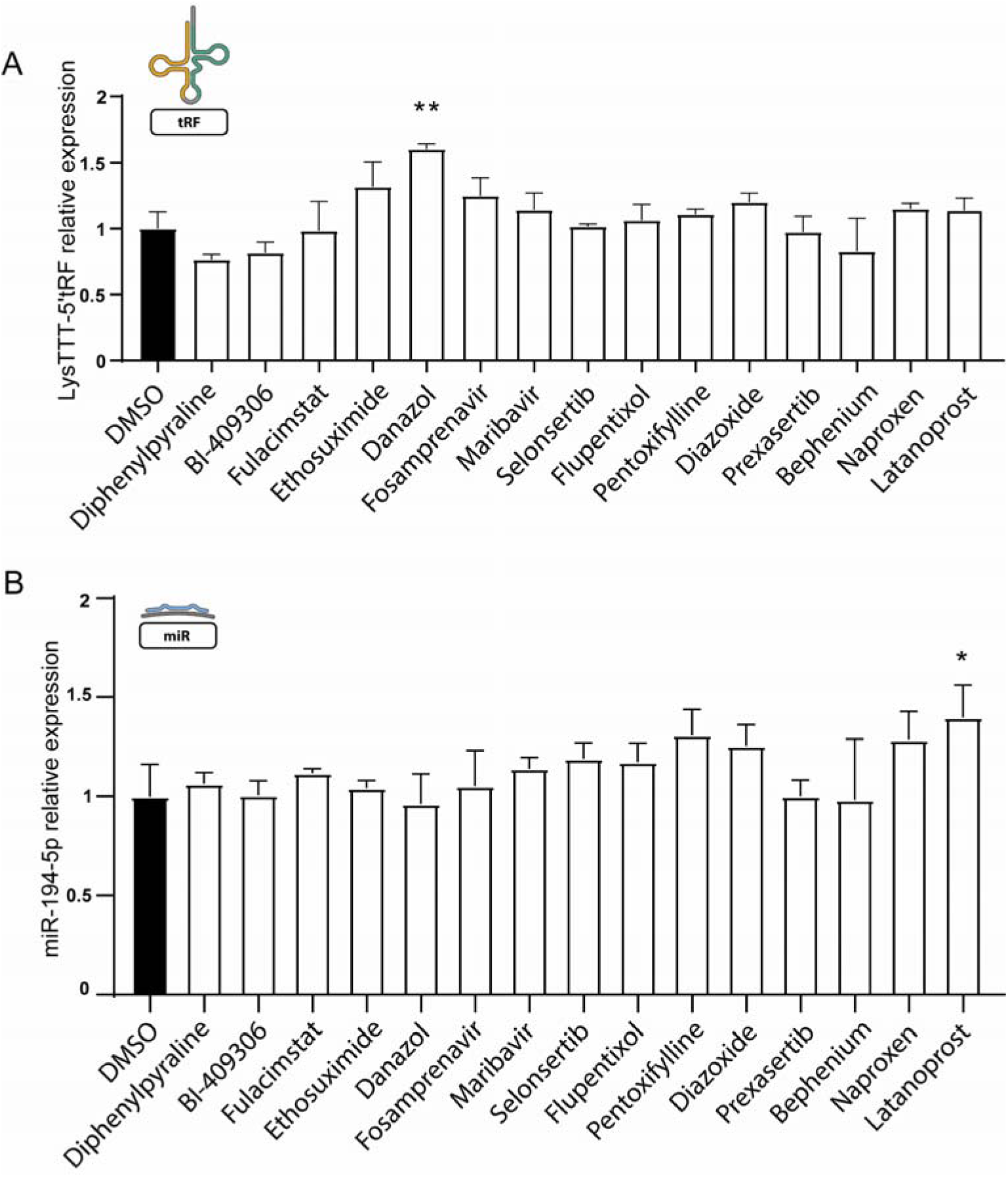
steatosis-alleviating Danazol and Latanoprost alter LysTTT-5’tRF and miR-194-5p levels. A. LysTTT-5’tRF levels are elevated following treatment with Danazol. tRF levels were measured by RT-qPCR after size selection and normalized to miR-145-5p and miR-15a-5p. B. miR-194-5p levels are elevated following treatment with Latanoprost. Measured by RT-qPCR and normalized to SNORD47. In all panels average ±SD, student’s t-test, * p <0.05, ** p <0.01, *** p <0.001.

**Supp. table 5:**
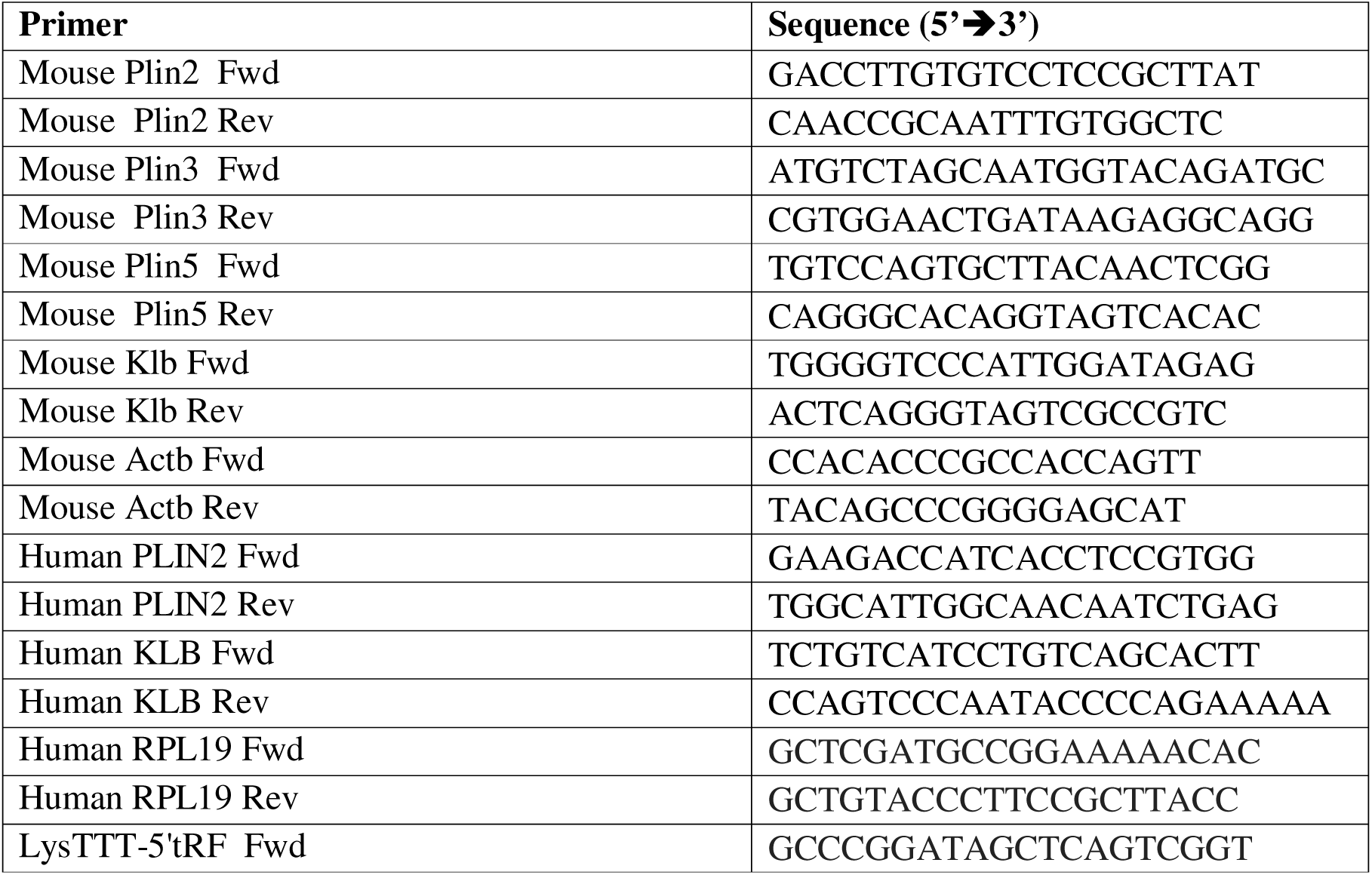
qPCR primers used in this research for quantification of mRNA targets and controls.

## Supplementary materials and methods

### Cell culture

Cells were grown at 37°C, 5% CO_2,_ in EMEM (Merck, M5650). Medium was supplemented with FCS (10% final concentration, Sartorius, 04-127), L-glutamine (2mM final concentration, Sartorius, 03-020) and Penicillin-Streptomycin-Amphotericin (100 units/mL, 0.10 mg/mL, 0.25 µg/mL, final concentrations, respectively, Sartorius, 03-033). Cells were mycoplasma free (EZ-PCR™ Mycoplasma Detection Kit, Sartorius). For experiments 150,000 cells/ml were plated and 24 hours later medium was replaced with oleic acid (OA)-rich medium prepared as described below. 24 hours after this cells were transfected with tRF mimic oligonucleotide or negative control (IDT, Syntezza Bioscience) using HiPerFect transfection reagent (Qiagen, 301705) and harvested after an additional 24 hours. Both oligos were 5’-phosphorylated RNA oligonucletotides with two 2’-O-methyl bases (labelled m), sequences as follows:

LysTTT-5’tRF mimic (25-mer): GC CCG GAU AGC UCA GUC GGU AGmA mGC.

Negative control (21-mer): GC GAC UAU ACG CGC AAU mAmUG.

### Oleic acid-rich medium

Full EMEM medium as above was supplemented with 2% BSA (Merck, A4919) and 0.5mM or 1mM OA (Merck, O1383). The medium was sonicated (Elmasonic S 30 (H), Elma) for 1 hour at 45°C to allow the formation of fatty acid-BSA complexes, stored overnight at 4°C and introduced to the cells.

### RT and qPCR

Synthesis of cDNA from mRNA and qPCR were done with qScript™ cDNA Synthesis Kit (Quantabio, 95047), PerfeCTa® SYBR® Green FastMix® (Quantabio, 95072), and human or mouse-specific primers (Suppl. Table 5). Synthesis of cDNA and qPCR for microRNA and tRFs were done with qScript™ microRNA cDNA Synthesis Kit (Quantabio, 95107), PerfeCTa® SYBR® Green FastMix® Low ROX (Quantabio, 95074) and human or mouse primers. Data is presented as relative normalized expression (ΔΔCt) with normalization as follows: all mRNAs to RPL19, all miRs to SNORD47, all tRFs to miR-15a-5p and miR-145-5p.

### Immunobloting

Cells were seeded in 12-well plates and incubated in 0.5 or 1mM OA-rich medium for 48 hours for PLIN2 quantification or 0.5mM OA-rich medium for 24 hours for KLB quantification, prior to transfection with LysTTT-5’tRF mimic for 24 hours as described above. For harvesting, cells were washed twice in cold PBS and incubated for 20 minutes on ice in RIPA buffer (50mM Tris-HCl pH 7.5, 150mM NaCl, 0.5% sodium deoxycholate, 1% Triton X-100, 0.1% SDS, 1mM EDTA, Protease Inhibitor Cocktail and Phosphatase Inhibitor Cocktail 1:100; Cell Signalling Technology, 5871 and 5870). Lysate was collected and centrifuged at 20,000 rpm at 4°C for 15 minutes, and supernatant transferred to fresh tubes. Protein concentrations were determined using Bradford (Merck, B6916) or Lowry (Bio-Rad, 5000113) assays and 5 µg/sample was loaded onto 4-15% gradient polyacrylamide gels (Bio-Rad, 4561083) and transferred (Bio-Rad, Trans-Blot Turbo Transfer System) to nitrocellulose membranes (Bio-Rad, 1704158). Proteins were detected with antibodies against PLIN2 (Proteintech, 15294-1-AP, 1:1500), KLB, (Abcam, ab106794, 1ug/ml) and Alpha-Tubulin (Merck, T5168, 1:1000), HRP-conjugated secondary antibodies (Jackson, 111-035-144, 115-035-062,), then ECL (Cell Signalling Technologies, 12757 or Thermo Fisher Scientific, 34580) using the myECL™ Imager, with its software for quantification (Thermo Fisher Scientific).

### Triglyceride quantification

#### Spectrophotometric quantification of lipid droplets

Cells were washed twice with ice-cold PBS, fixed for 30 minutes at room temperature in 4% PFA, washed twice with PBS, permeabilized by incubation for 10 minutes in 0.2% Triton X-100, and washed three times with PBS. Next, cells were incubated light-protected at RT in 1 µg/mL DAPI (Santa Cruz Biotechnology, sc-3598) for 5 minutes, washed three times with PBS, incubated in 300 nM Nile Red (Thermo Fisher Scientific, N1142) for 10 minutes and washed twice with PBS. A Tecan Spark™ 10M plate reader was used to quantify Nile Red signal (excitation 515 nm, emission 583 nm) which was then normalized to DAPI signal as indicator of cell number (exitation 370 nm, emission 440 nm).

#### Confocal image analysis

Quantification of lipid droplets was also carried out on cell images obtained by confocal microscopy (FLUOVIEW FV10i, Olympus). A total of ∼150 images for each condition were taken from multiple wells in three independent experiments (Nile Red, excitation 499 nm, emission 591 nm; DAPI, excitation 359 nm, emission 461 nm). Images were analyzed with ImageJ, with triglyceride content expressed as the total area of Nile Red signal normalized to the number of nuclei in the image. Detailed description of analysis: 3D images (oif. format) were split to Nile Red and nuclei channels and each was analyzed separately. For Nile Red, background was subtracted (rolling=30, sliding stack) and threshold calculated using all z-stack histograms. A 3D projection object with interpolation was created based on the post-threshold image. It was transformed into binary form and measured to extract the total area of signal. Nuclei were counted manually. To create colored pictures background was subtracted from the nuclei channel (rolling=10), merged with the Nile Red channel, and a z-stack projection object created using standard deviation of the different z-layers. The original Imagej Java code is submitted as a separate file.

### Size selection for tRF qPCR

1 ug total RNA from murine or Hep G2 samples were separated by PAGE (15% Criterion™ TBE-Urea Polyacrylamide Gel, Bio-Rad, 3450091). Bands containing RNAs less than 50 nucleotides long were excised from the gel based on markers (New England BioLabs, MicroRNA Marker, N2102S and Low Range ssRNA Ladder, N0364). To elute the RNA from the gel samples were rotated overnight at 4°C in 800 µL of 0.3M NaCl. 800 µL of isopropanol and 3 µL of glycogen (Merck, 10901393001) were added to each sample. After a 24-hour incubation at -20°C, RNA was precipitated by high-speed centrifugation (30 minutes, 20,000g, 4°C), washed in 1 ml of 70% ethanol, and recovered in 20 µL ddw. cDNA was synthesized from 500 pg and tRFs quantified with standard qPCR as described above (see Supp table 5 for primer sequences)

